# Multi-modal profiling of the extracellular matrix of human fallopian tubes and serous tubal Intraepithelial carcinomas

**DOI:** 10.1101/2021.09.21.461246

**Authors:** Carine Renner, Clarissa Gomez, Mike R Visetsouk, Isra Taha, Aisha Khan, Stephanie McGregor, Paul Weisman, Alexandra Naba, Kristyn S Masters, Pamela K Kreeger

## Abstract

Recent evidence supports the fimbriae of the fallopian tube as a potential origin site for high-grade serous ovarian cancer (HGSOC). The progression of many solid tumors is accompanied by changes in the microenvironment, including alterations of the extracellular matrix (ECM). The ECM of fallopian tube and HGSOC has not been well characterized. Therefore, we sought to determine the ECM composition of the benign fallopian tube and how it changes with the onset of serous intraepithelial carcinomas (STICs), precursor of HGSOC. The ECM composition of benign human fallopian tube was first defined from a meta-analysis of published proteomic datasets and identified 190 ECM proteins. We then conducted de novo proteomics using ECM enrichment and identified 88 proteins, 7 of which were not identified in prior studies. We further investigated the levels and localization of seven of these ECM proteins (type I, III, and IV collagens, fibronectin, laminin, versican, perlecan) and hyaluronic acid using multi-spectral immunohistochemical staining of fimbriae from patients with benign conditions or STICs. Quantification revealed an increase in stromal fibronectin and a decrease in epithelial versican in STICs. Our results provide an in-depth picture of the ECM in the benign fallopian tube and identified ECM changes that accompany STIC formation.

## INTRODUCTION

The most lethal and common of the epithelial ovarian cancers is high grade serous ovarian cancer (HGSOC), which is diagnosed in approximately 15,000 women annually in the United States. A majority of these patients will experience at least one recurrence following frontline treatment, resulting in a 5-year survival rate of only 48% [1]. A contributing factor to the high lethality of this cancer is its late-stage diagnosis when surgical debulking and chemotherapy, the primary means of treatment, may not be sufficient [2]. Diagnosis at late stages also obscured the cell of origin for the disease for decades; current literature now supports the ‘tubal hypothesis’, where the fallopian tube epithelium (rather than the ovarian surface epithelium) is the likely origin for many cases of HGSOC [3-5]. Support for the tubal hypothesis includes genomic analyses linking TP53 mutations in HGSOC samples to those found within the fallopian tube epithelium at sites known as serous tubal intraepithelial carcinomas (STICs) [6, 7]. Additional support for the tubal origin includes the stronger correlation of HGSOC tumors to the benign fallopian tube when comparing expression of the top 50 differentially expressed genes between potential origin sites of metastatic HGSOC (fallopian tube, ovary, and peritoneum, [5]).

While driver mutations of TP53 play an essential role in HGSOC, recent evidence also points to a role for the microenvironment, and more specifically of the extracellular matrix (ECM), in the progression of HGSOC. The ECM is a complex and dynamic meshwork of proteins that regulate cell and organ physiology and plays key roles in cancer initiation, progression, and dissemination [8, 9], including HGSOC [10]. For example, elevated levels of laminin-γ2, type I and type III collagens, fibronectin, versican, and hyaluronan have all been associated with a poorer prognosis of ovarian cancers [11]. Decreased type I collagen or increased type III collagen concentrations led to increased invasion of fallopian tube epithelial cells in models of ovarian cortical inclusion cysts, which are hypothesized to be the first metastatic site of HGSOC [12]. Evidence for ECM changes in STICs have been reported recently, including differences in collagen fiber morphology [13] and increased levels of highly sulfated chondroitin sulfate [14], an ECM component commonly associated with cancer [15].

While there is growing evidence to suggest that ECM changes contribute to the early progression of HGSOC, we still do not have a complete picture of the extent of ECM changes occurring as STICs arise in the fallopian tube. First, the ECM composition, or “matrisome”, of the benign fallopian tube has not been defined. A recent study characterized the changes in type I and III collagens and four proteoglycans in the follicular and luteal phases of the menstrual cycle [16]; however, this report focused on the ampullar region of the fallopian tube, where relatively few STICs have been observed [17]. Global proteomics of the human fallopian tube have been reported [18-21], but these methods are not able to differentiate intra- vs. extra-cellular components or characterize the insoluble matrisome in depth. An additional obstacle to analyzing ECM alterations in STICs is the limited availability of STIC samples from pre- or early stage HGSOC. STICs are primarily identified when patients undergo surgery for suspected ovarian cancer; at this point, the tumor has frequently advanced to stage III with peritoneal metastases, and STICs may be altered by the systemic changes associated with increased tumor burden. Alternatively, STICs in pre- or early-stage HGSOC can be identified during risk-reduction salpingectomy followed by sectioning and extensively examining the fimbria (SEE-FIM) [3]. Moreover, when STICs are identified at these early stages, they are restricted regions of tens to hundreds of cells, which limits the type and number of analyses that can be performed. Recent work employed a combination of laser capture microdissection and ultra-high-sensitivity mass spectrometry to perform proteomic profiling of STICs from patients with metastatic disease [18]. Although the generated STIC proteomic signature contained many ECM proteins, the method employed was not ideal for capturing the insoluble ECM, suggesting that it did not completely define the tissue matrisome.

Here, we employed a set of complementary approaches to characterize the ECM of human fallopian tubes and examine how this ECM changes when STIC lesions develop. We first conducted a meta-analysis of existing global proteomic profiles of benign human fallopian tube studies to identify the ECM and ECM-associated proteins found in these tissues. We next used an ECM-enrichment approach to characterize by mass spectrometry the insoluble ECM components composing the human fallopian tube matrisome. Based on the result of our experimental ECM data and meta-analysis, we then utilized multispectral immunofluorescence to characterize the differences in a selected set of core matrisome components including type I, III, and IV collagens, the proteoglycans perlecan and versican, the glycoproteins fibronectin and laminins, and the glycosaminoglycan hyaluronan, in STICs compared to non-STIC regions of patients with STICs or to patients with benign fallopian tubes. Elucidating the changes in the ECM during STIC development could provide critical insights into the pathogenic progression of HGSOC, and potentially illuminate diagnostic markers and preventative therapeutic strategies.

## MATERIALS AND METHODS

Unless specified otherwise, all reagents were from ThermoFisher Scientific, Waltham, MA, USA.

### Meta-analysis of existing proteomic studies of human fallopian tubes

Global proteomic abundance data were collected from the supplementary tables of four studies: Wang et al., Table S1 [19]; Eckert et al., Table S2 [18]; Hu et al., Table S2 [20]; and McDermott et al., Table S1 [21]. Data for benign [19] or histologically normal [18, 20, 21] human fallopian tube (hFT) samples were extracted from each study, representing data from >100 unique patient samples, and data for STIC stromal and STIC epithelial samples were extracted from the study by Eckert et al. Proteins detected with nonzero abundance in any single sample within a given study were added to that study’s hFT protein list. Each list was then annotated to identify matrisome proteins, and matrisome proteins were further classified into the different matrisome divisions and categories we previously defined [22, 23] (see Supplementary Table 1).

### Proteomic analysis of human fallopian tube

#### Tissue decellularization

Four flash-frozen benign human fallopian tube samples from four Caucasian women individuals (age 40-52) were procured from the University of Wisconsin Carbone Cancer Center Translational Science Biocore BioBank, which collects biospecimens through an IRB-approved protocol at UW-Madison and then acts as an honest broker to provide deidentified tissue to investigators. Since we obtained a large sample (700 mg) from one of the individuals, we further divided this sample into three 150 mg samples, allowing us to assess tissue heterogeneity.

100-150 mg of tissue were homogenized for 2 minutes at a power setting of 12 using a Bullet Blender and stainless-steel beads of varying diameters (Navy bead lysis kit, Next Advance, Averill Park, NY) following manufacturer’s instructions. ECM enrichment was achieved through the sequential extraction of intracellular proteins in order of decreasing protein solubility, using a subcellular protein fractionation kit for tissues following the manufacturer’s instructions. The efficiency of the sequential extraction of intracellular components and concomitant ECM-protein enrichment was monitored by western blot analysis, probing for collagen I (Sigma, #AB765P, 1 μg/mL, St. Louis, MO), histone H4 (Sigma, #05-858, 1 μg/mL), and actin using a serum containing anti-actin antibodies (1/5000) kindly gifted by the Hynes lab at Massachusetts Institute of Technology, Cambridge, MA, USA.

#### Protein sample preparation for mass spectrometry

The ECM-enriched protein sample was subsequently solubilized and digested into peptides following an established protocol [22, 24]. Briefly, proteins were solubilized in an 8 M urea solution prepared in 100 mM NH_4_HCO_3_, and protein disulfide bonds were reduced using 10 mM dithiothreitol. Reduced disulfide bonds were then alkylated with 25 mM iodoacetamide (30-minute incubation, in the dark, at room temperature). The urea concentration was brought to 2 M, and proteins were then deglycosylated with PNGaseF (New England Biolabs, Ipswich, MA) for 2 hours at 37C, in Celsius units and digested sequentially, first with Lys-C for 2 hours at 37C, in Celsius units, and then with trypsin, overnight at 37C, in Celsius units. A fresh aliquot of trypsin was added on the following day and samples were incubated for an additional 2 hours at 37C, in Celsius units. All incubations were performed under agitation as previously described. The sample was acidified with 50% trifluoroacetic acid (TFA) and desalted using Pierce Peptide Desalting Spin Columns. Peptides were reconstituted in 95% HPLC grade water, 5% acetonitrile and 0.1% formic acid, and the concentration of the peptide solution was measured using the Pierce Quantitative Colorimetric Peptide Assay kit.

#### Peptide analysis by LC-MS/MS

Approximately 600 ng of desalted peptides were analyzed at the University of Illinois Mass Spectrometry core facility on a quadrupole Orbitrap mass spectrometer Q Exactive HF mass spectrometer coupled with an UltiMate 3000 RSLC nano system with a Nanospray Frex Ion Source (Thermo Fisher Scientific). Three technical replicates (hFT1-1, hFT1-2, hFT1-3) were acquired for one patient sample and one technical replicate for each of the other three patient samples. Samples were loaded into a PepMap C18 cartridge (0.3 × 5mm, 5μm particle) trap column and then a 75 μm x 150 mm PepMap C18 analytical column (Thermo Fisher Scientific) and separated at a flow rate of 300 nL/min. Solvent A was 0.1% formic acid (FA) in water and solvent B was 0.1% FA, 80% acetonitrile (ACN) in water. The solvent gradient of LC was 5–8% B in 5 min, 8-10% B in 7 min, 10-30% B in 65 min, 30-40% B in 80 min, 40-95% B in 90 min, wash 95% B in 95 min, followed by 5% B equilibration until 105 min. Full MS scans were acquired in the Q-Exactive mass spectrometer over 350-1400 m/z range with resolution of 60,000 (at 400 m/z) from 5 min to 97 min. The AGC target value was 1.00E+06 for full scan. The 15 most intense peaks with charge state 2, 3, 4, 5, 6 were fragmented in the HCD collision cell with normalized collision energy of 30%, these peaks were then excluded for 12s within a mass window of 1.2 m/z. Tandem mass spectrum was acquired in the mass analyzer with a resolution of 15,000. The AGC target value was 5.00E+04. The ion selection threshold was 1.00E+04 counts, and the maximum allowed ion injection time was 30 ms for full scans and 50 ms for fragment ion scans.

#### Database searching

All MS/MS samples were analyzed using Mascot (Matrix Science, London, UK; version 2.6.2). Mascot was set up to search the Uniprot-human_20200811 database (unknown version, 20375 entries) assuming the digestion enzyme stricttrypsin. Mascot was searched with a fragment ion mass tolerance of 0.20 Da and a parent ion tolerance of 10.0 PPM. O-110 of pyrrolysine, u+49 of selenocysteine and carbamidomethyl of cysteine were specified in Mascot as fixed modifications. Gln->pyro-Glu of the n-terminus, deamidated of asparagine and glutamine and oxidation of methionine, lysine, and proline were specified in Mascot as variable modifications, the latter two being characteristic post-translational modifications of ECM proteins, in particular collagens and collagen-domain-containing proteins, as we previously reported [25]. Of the six samples processed, only the three that came from the same individual, a 43-year-old Caucasian female diagnosed with a benign pelvic mass, had a quality sufficient for subsequent analysis.

#### Criteria for protein identification

Scaffold (version Scaffold_4.11.1, Proteome Software Inc., Portland, OR) was used to validate MS/MS based peptide and protein identifications. Peptide identifications were accepted if they could be established at greater than 91.0% probability to achieve a FDR less than 1.0% by the Scaffold Local FDR algorithm. Protein identifications were accepted if they could be established at greater than 8.0% probability to achieve an FDR less than 1.0% and contained at least 2 identified peptides. Protein probabilities were assigned by the Protein Prophet algorithm [26]. Proteins that contained similar peptides and could not be differentiated based on MS/MS analysis alone were grouped to satisfy the principles of parsimony. Proteins sharing significant peptide evidence were grouped into clusters. Mass spectrometry output was further annotated to identify ECM and non-ECM components [22, 25]. Specifically, matrisome components are classified as core-matrisome or matrisome-associated components, and further categorized into groups based on structural or functional features: ECM glycoproteins, collagens, or proteoglycans for core matrisome components; and ECM-affiliated proteins, ECM regulators, or secreted factors for matrisome-associated components (Supplementary Table 2, [22, 23]).

Raw mass spectrometry data have been deposited to the ProteomeXchange Consortium [27] via the PRIDE partner repository [28] with the dataset identifier PXD023707. The raw data will be made publicly available upon acceptance of the manuscript.

### Multispectral analysis of human fallopian tubes and STICs

#### Tissue procurement

Archived formalin-fixed, paraffin-embedded fallopian tube/fimbriae (10 benign, 12 with STICs) were acquired from archived pathology samples through an IRB-approved protocol at UW-Madison. Samples were retrieved from cases of benign or STIC patients that had undergone surgical debulking. Clinical information can be found in Supplementary Table 3.

#### Multispectral immunohistochemistry

Formalin-fixed, paraffin-embedded fallopian tube/fimbriae were cut and processed into 5 μm sections as previously described [29]. Multispectral immunohistochemistry was performed using the Opal 7-Color Manual IHC Kit (Akoya Biosciences/PerkinElmer, NEL811001KT, Marlborough, MA) according to the manufacturer’s protocol. The maximum number of fluorescent signals that could be spectrally unmixed was limited to 5 channels; therefore, we separated our analyses of these ECM components, along with nuclei and P53 for tissue context and STIC identification, into two staining sets (Supplementary Table 4). Briefly, slides were deparaffinized using SafeClear II xylene substitute followed by rehydration in a series of ethanol dilutions (100% twice, 90%, 70%, 50%) ending with a rinse in deionized water. Slides were fixed in 10% neutral buffered formalin for 20 minutes, rinsed in deionized water and then placed in PerkinElmer AR6 buffer. For antigen retrieval, slides were microwaved at 100% in AR6 buffer until the buffer boiled and then microwaved at 20% power for 15 minutes, cooled to room temperature, and rinsed in deionized water and tris-buffered saline with 0.1% tween-20 (TBST). PerkinElmer blocking solution was used overnight at 4C, in Celsius units, followed by biotin blocking with avidin (Vector Laboratories, SP-2001, Burlingame, CA) for 15 minutes, TBST wash, and then biotin incubation for 15 minutes. Optimized antibody information is in Supplementary Table 4. Briefly, set 1 consisted of type IV collagen, pan-laminin, type I collagen, type III collagen, and P53, stained in that order. Set 2 consisted of hyaluronan, perlecan, fibronectin, versican, and P53, stained in that order. Slides were incubated in primary antibody solutions for two hours at room temperature, washed in TBST, then incubated in PerkinElmer Polymer HRP Ms+Rb for ten minutes at room temperature. Slides were washed in TBST and incubated in Opal Fluorophore diluent as described in Supplementary Table 4 for ten minutes. Antibody stripping was performed in AR6 buffer in between each component using the microwave technique described for antigen retrieval. Slides were blocked for thirty minutes at room temperature or overnight at 4C, in Celsius units, followed by avidin and biotin blocking, and the staining process was repeated. After all ECM components and P53 had been stained, slides were counterstained with DAPI and sealed using ProLong Diamond Antifade Mountant.

#### Imaging and Analysis

Imaging of fimbriae was performed on a Nuance Multispectral imaging system with an Olympus UPlanFL N 20x objective, NA 0.5, using the Nuance Software version 3.0.2 (PerkinElmer). Spectral unmixing of the Opal fluorophores and autofluorescence was performed post-imaging using manufacturer-provided instructions, and images were exported as 16-bit TIFF. Images were categorized as either benign or STIC, based on patient source. All non-STIC patients had single-cell layered epithelia with only occasional P53-positive cells. Within STIC patients, we observed both STIC regions (three or more layers of P53-positive cells) and non-STIC regions (single-cell layered epithelia, P53-negative), (Supplementary Figure 1). As samples were from pathology archives, it is possible that the entire STIC was sectioned during clinical analysis. Indeed, not all sections from STIC patients had STIC regions, but were included in non-STIC region analyses based on the associated pathological report. Regions of interest (ROIs) were created for epithelia or stroma using the DAPI image, free-hand selection tool, and ROI manager in FIJI ImageJ Software version 2.0.0 (Supplementary Figure 2). Images were analyzed for mean fluorescent signal intensity (MFI) in epithelial and stromal ROIs, and their relative levels calculated as an epithelial to stromal ratio (E:S ratio).

#### Statistical Analysis

Statistical calculations were carried out as indicated in the figure legends (GraphPad Prism v8.4.1) with p<0.05 set as a threshold for significance.

## RESULTS

### Meta-analysis of the proteomes of benign human fallopian tube and STICs

Although the ECM of the fallopian tube has not yet been fully characterized, the global proteome of the benign and HGSOC fallopian tubes has been profiled in several recent studies [18-21]. We thus sought to conduct a meta-analysis of these studies with the goal of defining the human fallopian tube matrisome. To do so, we retrieved the list of proteins identified in human fallopian tubes or STIC samples profiled in the four studies representing data from >100 unique patient samples [18-21] (Supplementary Table 1A) and annotated them to determine which proteins belong to the human matrisome that we previously defined [22]. This led to the identification of 295 ECM proteins in the Wang dataset, 238 in the Eckert dataset, 397 in the Hu dataset, and 524 in the McDermott dataset (Fig. 1A, Supplementary Table 1B). These vastly different numbers can be explained by multiple factors, including the depth (i.e., were the peptides fractionated or not) and modality of the mass spectrometry analyses, and the number of samples included in each study (Supplementary Table 1A). We next broke down the proteins into different matrisome categories. First, there are the structural ECM components that constitute the “core matrisome”: collagens, proteoglycans and ECM glycoproteins. In addition, we identified “matrisome associated” proteins, including ECM-associated proteins, ECM regulators, and secreted factors. In all four studies, the majority of matrisome proteins detected belonged to the matrisome-associated class, including ECM regulators that are enzymes known to impact ECM proteins, such as proteases (Fig. 1A, Supplementary Table 1B). The comparison of the data from these four studies led to the identification of a set of 190 matrisome proteins present in all four human fallopian tube proteome datasets out of initial 430 distinct matrisome proteins identified across the individual studies (Fig. 1B, Supplementary Table 1B).

**Figure 1.**
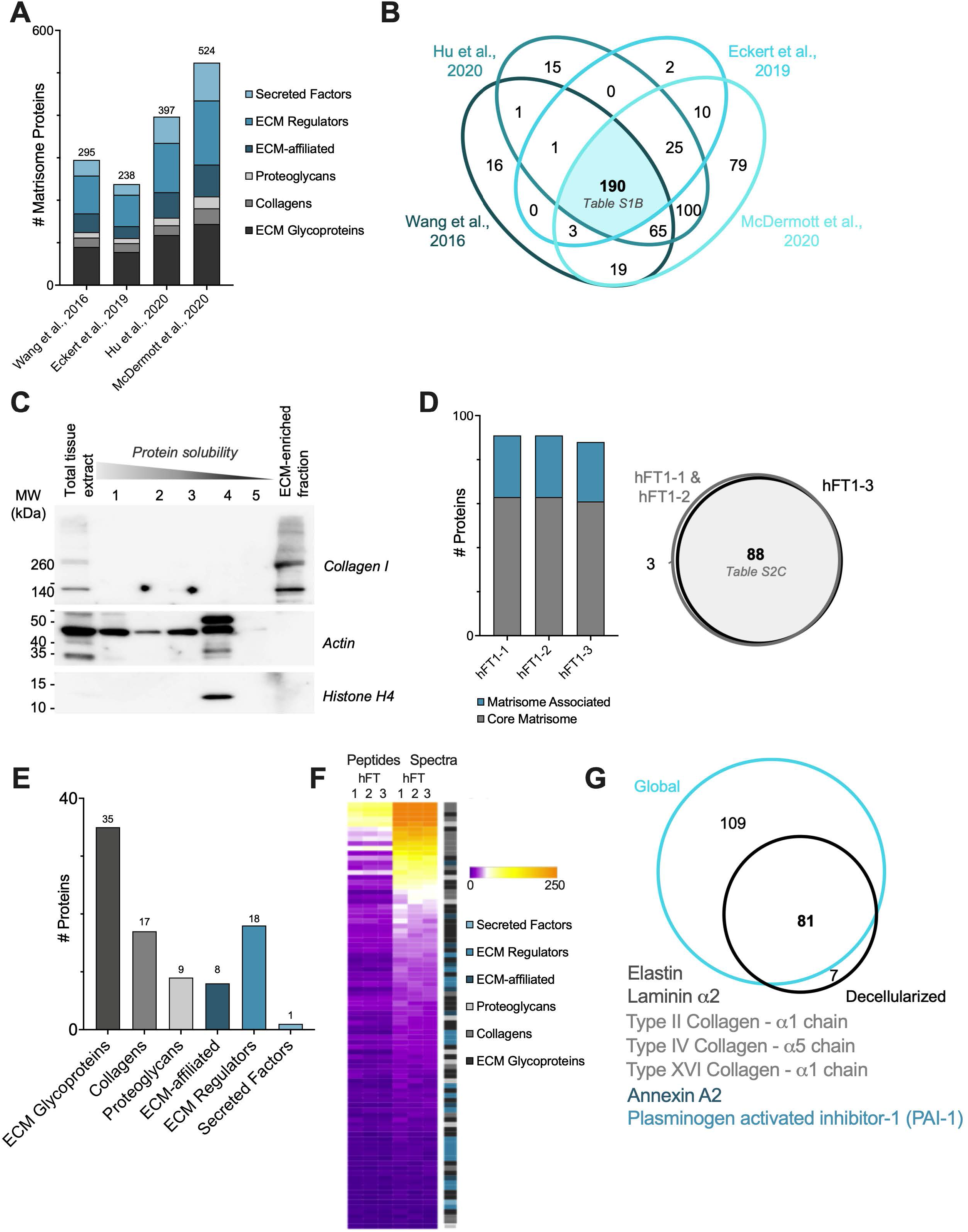
Proteomic analysis of the human fallopian tube matrisome. (A) Distribution of ECM and ECM-associated proteins detected in benign human fallopian tube samples in four global proteomic studies (see Supplementary Table 1B). (B) Venn diagram of matrisome proteins detected in four global proteomic studies (see Supplementary Table 1B). (C) Western blot analysis of different protein fractions obtained during sample decellularization process revealed successful solubilization and extraction of intracellular proteins (actin, histones) while the diagnostic ECM protein, type I collagen, was retained in the remaining insoluble fraction. The ECM-enriched protein fraction was further analyzed by mass spectrometry. (D) The three technical replicates (hFT1-1, hFT1-2, hFT1-3) resulted in a similar number of matrisome proteins (left panel), with a strong degree of overlap in identity (right panel, see Supplementary Tables 2B and 2C). (E) Distribution of ECM and ECM-associated proteins detected among matrisome categories (see Supplementary Table 2C). (F) Heatmap representing normalized numbers of unique peptides and total spectra detected for each matrisome protein in all three replicates. (G) Venn diagram of the experimental matrisomic data obtained in comparison to the meta-analysis of global proteomic studies of human fallopian tubes. Unique proteins are listed, font color indicates matrisome category.

One of the problems inherent in the proteomic analysis of STIC tissues is the very small sample size. To overcome this, Eckert and collaborators [18] developed a novel method that combined laser capture microdissection and label-free MS to identify proteins present in samples composed of as few as 5,000 cells. This method also allowed the tumor compartment to be separated from the stromal compartment, with the latter presumably enriched for ECM proteins. However, the tissue used for this proteomic analysis was collected from patients who had already progressed to metastatic HGSOC; in contrast, we seek to characterize the earliest stages using benign fallopian tubes and STIC lesions in the absence of metastatic tumors.

### Proteomic analysis of benign human fallopian tube/fimbriae

Global proteomics permits the identification of a significant number of ECM and ECM-associated proteins, but whether these proteins are, in fact, part of the assembled ECM meshwork remains to be determined. Indeed, while all ECM and ECM-associated proteins are produced intracellularly, they exert their functions when they are incorporated in the insoluble ECM scaffold. The incorporation of the proteins in the ECM scaffold requires post-translational modifications, including glycosylation and cross-linking, and protein-protein interactions. These factors render ECM and ECM-associated proteins largely insoluble and so the pool of assembled ECM proteins is notoriously difficult to capture from pre-cleared tissue lysates [25, 30], which were utilized in the studies included in our meta-analysis.

We previously developed a mass-spectrometry pipeline tailored to study insoluble ECM proteins [22, 24]. In brief, this pipeline consists of 1) enrichment of ECM proteins from tissue and concomitant depletion of soluble intracellular components, 2) digestion of ECM-enriched protein samples into peptides after deglycosylation to maximize trypsin accessibility and proteolytic cleavage, 3) acquisition of mass spectrometry data, and 4) analysis of mass spectrometry data, with the inclusion of parameters to maximize ECM protein identification, such as allowing for ECM-specific post-translational modifications such as hydroxylysines and hydroxyprolines [25, 30]. Here, to experimentally characterize the protein composition of the insoluble ECM fraction of fallopian tube/fimbriae, we obtained samples from patients without a history of ovarian cancer and enriched the ECM as previously described [24]. This enrichment was monitored by conducting western blots on all protein fractions generated during the decellularization and probing for collagen I, cellular actin, and nuclear histone H4 (Fig. 1C). While we were able to enrich for ECM proteins from four independent samples and conduct preliminary MS/MS analysis of each (data not shown), only one sample generated enough material for a robust MS/MS analysis. Identification of the proteins from the ECM-enriched fraction from this sample was performed through mass spectrometric analysis. To determine technical variability in the list of proteins identified, 3 technical replicates (hFT1-1, hFT1-2, hFT1-3) were performed and revealed a high level of reproducibility (Fig. 1D, Supplementary Figure 3, Supplementary Table 2B). Overall, we consistently identified 88 matrisome proteins in the ECM of the human fallopian tube sample, including 35 glycoproteins, 17 collagens, and nine proteoglycans in the category of core matrisome components; and nine ECM-affiliated proteins, 18 ECM regulators, and one secreted factor in the category of matrisome-associated proteins (Fig. 1E, Supplementary Table 2C). Each of these 88 proteins were also identified in at least one of the other samples that had only preliminary analysis conducted (data not shown). Therefore, we proceeded with analysis of the draft fallopian tube matrisome. Using normalized spectral counts to estimate relative protein abundance, we found that the most abundant proteins were core matrisome proteins (Fig. 1F, Supplementary Table 2C).

Last, we sought to compare our findings to that of the meta-analysis we conducted on global proteomic datasets. While the scale and depth of our study is very small in comparison to that of the previously published studies, we were able to detect seven proteins that were not detected by global proteomics. These include three collagens: the fibrillar type II collagen (COL2A1), the basement membrane collagen COL4A5, and type XVI collagen (COL16A1) of the fibril-associated collagens with interrupted helices (FACIT) collagen family (Fig. 1G, Supplementary Tables 1B and 2C). Two other core ECM glycoproteins were detected: elastin and the α chain of laminin 5, as well as annexin A2 and the plasminogen activator inhibitor-1 (PAI1 or SERPINE1). In addition to many other proteins, we detected subunits of type I collagen, type III collagen, type IV collagen, and laminins; as well as fibronectin (FN1), versican (VCAN), and perlecan (HSPG2) through both our findings and the previous meta-analysis (Supplementary Tables 1B and 2C). Of these, collagen I, collagen III, fibronectin, versican, and laminin have previously been associated with poorer prognosis in ovarian cancer [11]. Again, each of these proteins was identified in at least one of the other fallopian tube samples that was only able to be preliminarily analyzed. Altogether, our pilot analysis demonstrated the feasibility of applying ECM-focused proteomics to gain insight into the composition of the ECM scaffold of human fallopian tube.

### Characterization of the ECM of STICs using multispectral immunofluorescence

As noted above, early-stage STIC samples are rare due to the high frequency of late-stage diagnosis of HGSOC. Moreover, common ECM analysis techniques, such as the mass spectrometry pipeline described above or even immunohistochemical staining of numerous tissue slices, require more material than is typically present in an early STIC. Thus, ECM changes that occur during STIC development remain largely unexplored. Building upon the human fallopian tube matrisome identified in Fig. 1, we aimed to investigate differences in the ECM of STICs versus benign fallopian tube tissue using multispectral immunofluorescence, which enables multiplexed visualization of ECM components (Fig. 2, Supplementary Figure 4), and is thus advantageous for scarce samples. For subsequent analysis, we selected a panel of seven core matrisome proteins that were detected with high confidence in our proteomic analysis of the ECM of benign fallopian tube as well as in the global proteomic studies of human fallopian tubes (see above). The panel comprises the fibrillar type I and III collagens, the basement membrane type IV collagen, the basement membrane proteoglycan perlecan (HSPG2), and the proteoglycan versican (VCAN), pan-laminin, and fibronectin. We also analyzed levels of the glycosaminoglycan hyaluronan (HA), based on prior reports of HA upregulation in the cancer microenvironment and role in cancer progression [31, 32]. In addition to analyzing differences in the ECM composition between benign and STIC patients, we examined differences in ECM levels in STIC regions versus non-STIC regions within the same patient (Supplementary Figure 1). Despite the relative rarity of early STIC samples, we were able to profile each target in a minimum of seven unique patients for each category (benign, non-STIC, and STIC regions).

**Figure 2.**
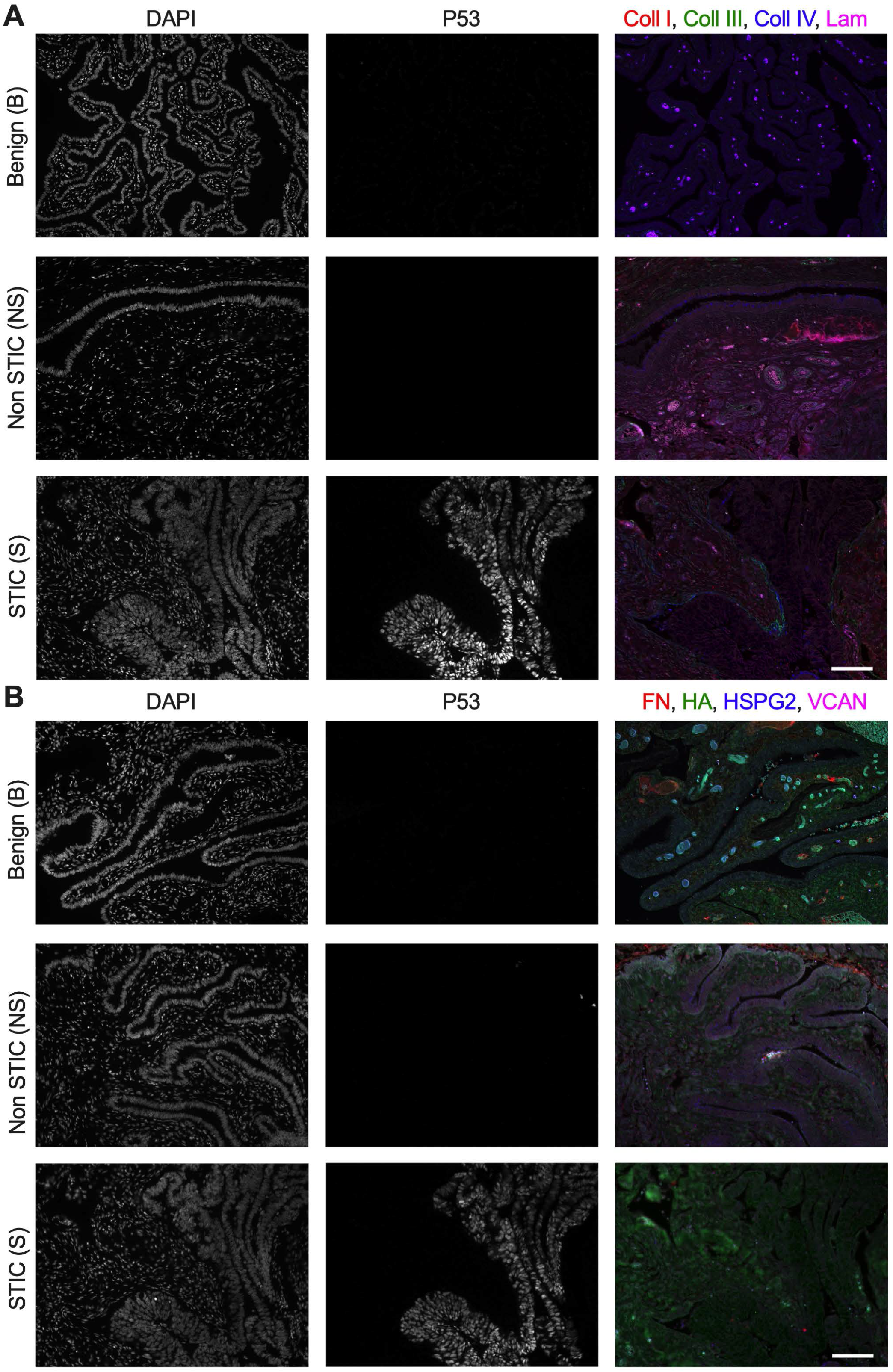
Differential levels of ECM in fimbriae of benign versus STIC patients. (A) Representative multispectral staining for DNA (DAPI), P53, and ECM components: type I collagen (Coll I, red), type III collagen (Coll III, green), type IV collagen (Coll IV, blue), pan-laminin (Lam, magenta) from a benign sample, and non-STIC and STIC region from a STIC patient. (B) Representative multispectral staining for DNA (DAPI), P53, and ECM components: fibronectin (FN, red), hyaluronan (HA, green), perlecan (HSPG2, blue), versican (VCAN, magenta) from a benign sample, and non-STIC and STIC region from a STIC patient. Scale bar = 100 μm.

### Distribution of collagens in STIC ECM

Examination of staining for type I, type III, and type IV collagens indicated that these proteins were present in both the epithelial and stromal regions of benign fallopian tube/fimbriae, non-STIC regions, and STICs (Fig. 2A). To define the ECM changes occurring in the epithelium and in the stroma immediately adjacent to the epithelium, we quantified the mean fluorescent intensity (MFI) of the multispectral ECM staining in selected regions of interest (Supplementary Figure 2). Comparisons were made between benign and non-STIC tissue, benign and STIC tissue (Fig. 3A), and between patient-matched non-STIC and STIC regions (Supplementary Figure 5). In both the epithelial and stromal regions, the intensity of each ECM protein was similar for all comparisons (Fig. 3B,C). We noted that there was substantial variability in the levels of type IV collagen in STIC patients. Comparison of non-STIC to STIC regions within the same patient showed a non-significant trend of decreased type IV collagen in STICs (Supplementary Figure 5). We next examined the potential for changes in ECM localization by calculating the ratio of the epithelial to stromal signal for each protein. Consistent with other tissues [33-35], the E:S ratio of benign tissue indicated that type I collagen was more abundant in the stroma (E:S less than 1.0), while type III and type IV collagens were distributed nearly evenly between the epithelia and stromal regions (E:S approximately 1.0). Interestingly, the E:S ratios of type III and type IV collagens were significantly decreased in STICs relative to benign tissue (Fig. 3D), and showed a similar, but not significant, trend of decreasing between patient-matched STIC and non-STIC regions (Supplementary Figure 5). Given the limited number of matched samples, it is difficult to be certain if the change in E:S resulted from loss of epithelial type III and type IV collagens or an increase in stromal type III and type IV collagens; regardless, these results suggest a relative enrichment in stromal collagens compared to benign or non-STIC regions.

**Figure 3.**
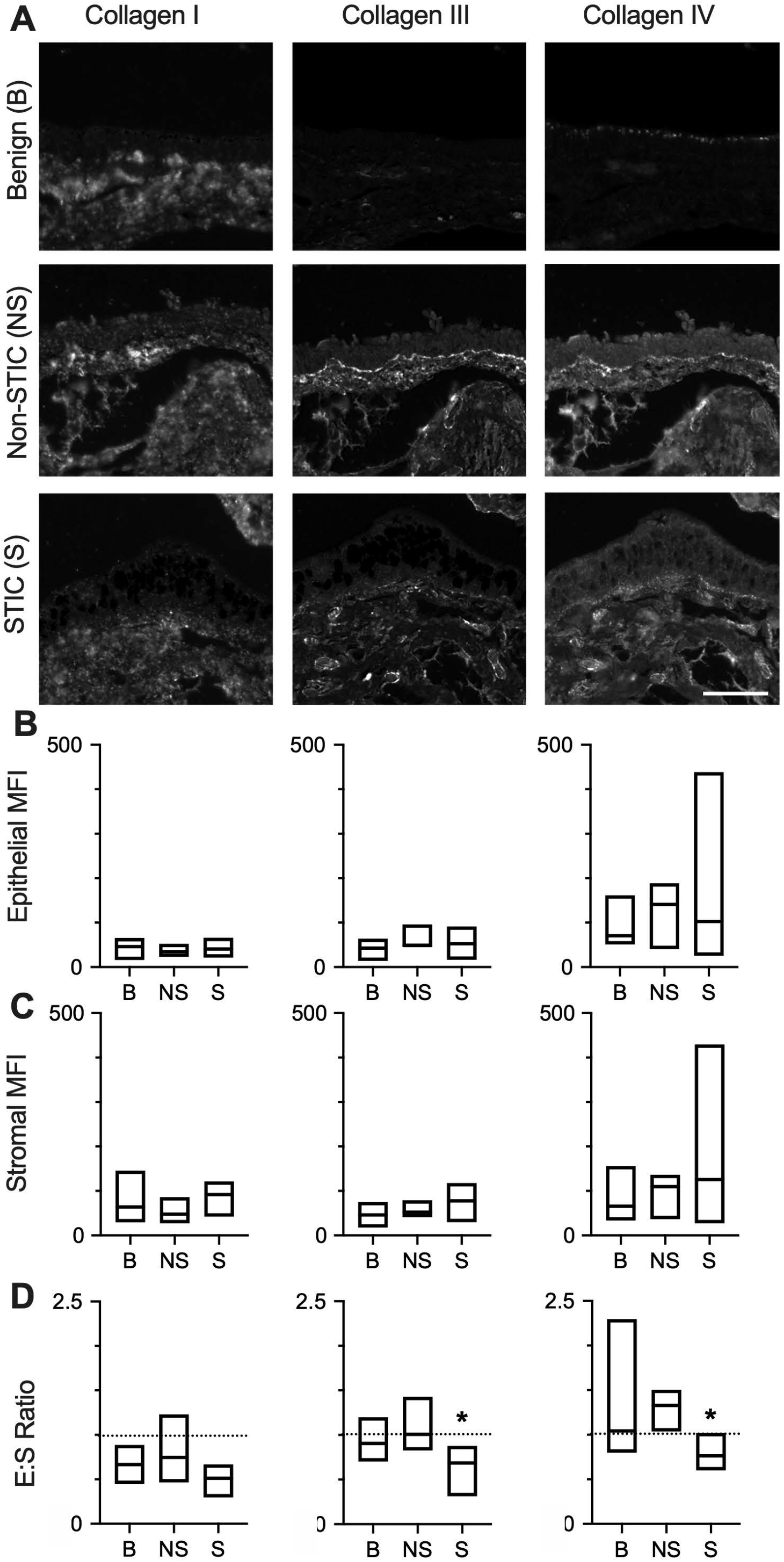
Differential levels of collagens in STICs. (A) Representative multispectral staining of fimbriae for type I, type III, and type IV collagens in benign (B), non-STIC (NS), or STIC (S) regions. (B-D) Quantification of (B) epithelial MFI, (C) stromal MFI, and (D) epithelial:stromal ratio (E:S ratio) of collagen I, collagen III, or collagen IV in B, NS, and S regions. Data are presented as a box plot indicating median and range, dashed line demonstrates E:S = 1. * indicates p< 0.05 for S compared to B by Kruskal-Wallis test and Dunn’s multiple comparison post-test. n=10 (B), 8 (NS), 7 (S). Scale bar = 200 μm.

### Distribution of proteoglycans and GAGs in the STIC ECM

Multispectral staining indicated the presence of hyaluronan, perlecan, and versican in epithelial and stromal regions of benign fallopian tube/fimbriae, non-STIC regions, and STICs (Fig. 2B). The levels of hyaluronan and perlecan were not significantly different for any direct comparisons of epithelial and stromal regions (Fig. 4A-C). While versican did not show a difference between regions at a population level, there was a significant decrease in epithelial versican between patient-matched STIC regions relative to non-STIC regions (Supplementary Figure 5). In benign tissue, analysis of the E:S ratio showed that hyaluronan and perlecan levels were both similar across the epithelia and stroma, while versican was more abundant in the epithelia compared to the stroma (Fig. 4D). Although the E:S ratio for hyaluronan was consistent across the three tissue types, perlecan and versican exhibited lower E:S ratios in STICs compared to benign tissue (Fig. 4D), and in STICs compared to non-STIC regions of patient-matched tissues (Supplementary Figure 5). For perlecan, examination of the individual samples suggests this is due to loss of epithelial production, as all but one patient had a decrease in epithelial perlecan (Supplementary Figure 5; note that the outlier had increases in both epithelial and stromal perlecan but an overall drop in the E:S ratio). As noted above, the decrease in epithelial versican was statistically significant between matched samples, confirming that the decreased E:S ratio resulted from relative depletion of versican in the epithelial layer.

**Figure 4.**
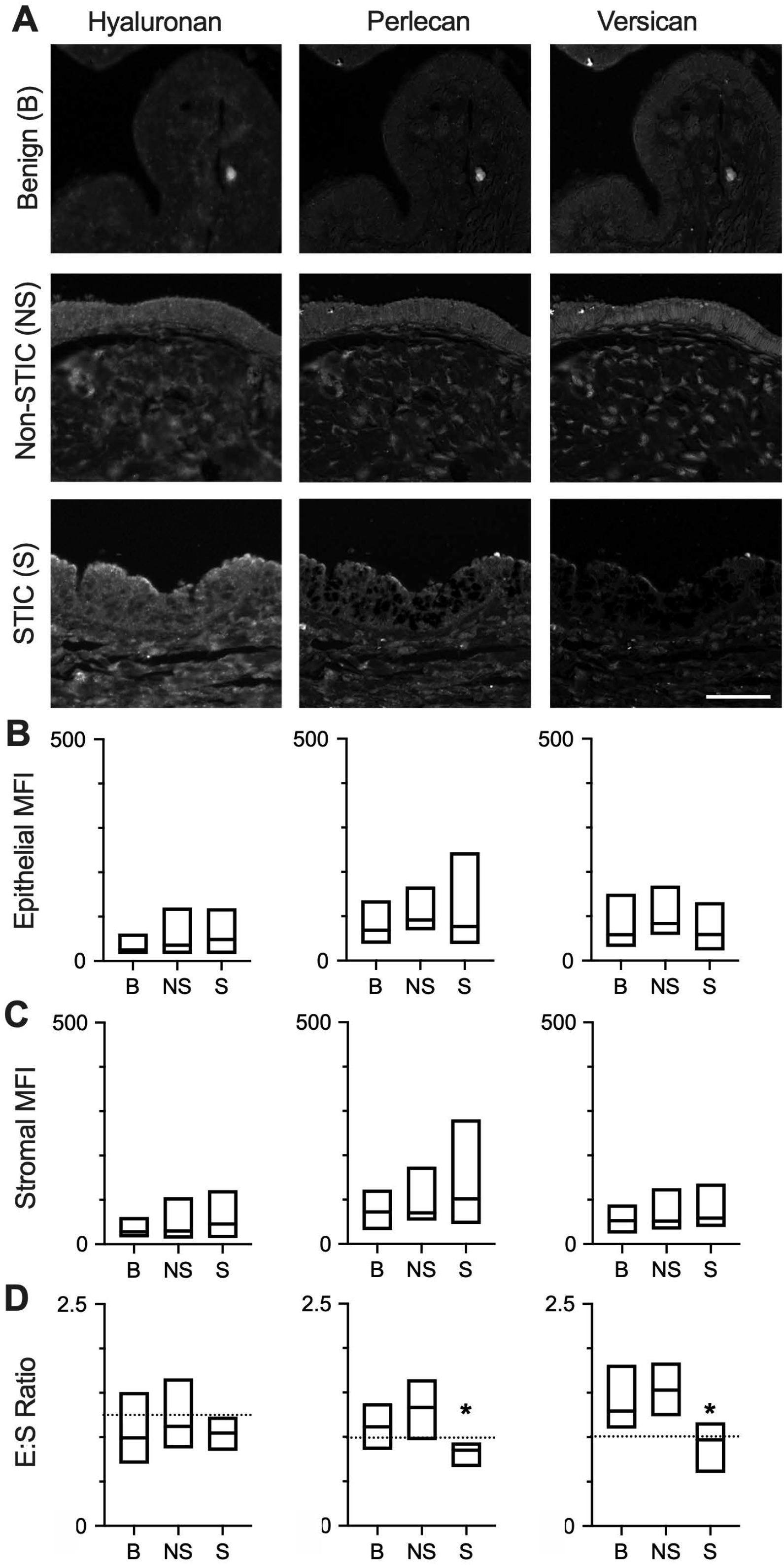
Differential levels of proteoglycans and GAGs in STICs. (A) Representative multispectral staining of hyaluronan, perlecan, or versican in benign (B), non-STIC (NS), and STIC (S) regions. (B-D) Quantification of (B) epithelial MFI, (C) stromal MFI, and (D) epithelial:stromal ratio (E:S ratio) of hyaluronan, perlecan, or versican in B, NS, or S regions. Data are presented as box plots indicating median and range, dashed line demonstrates E:S = 1. * indicates p< 0.05 for S compared to B by Kruskal-Wallis test and Dunn’s multiple comparison post-test. n=10 (B), 11 (NS), 8 (S). Scale bar = 200 μm.

### Distribution of glycoproteins in STIC ECM

Laminin was observed in multispectral staining of epithelial and stromal regions of benign fallopian tube/fimbriae, non-STIC regions, and STICs (Fig. 2A). Laminin in epithelial and stromal regions was not different between all comparisons (Fig. 5A-C, Supplementary Figure 5). The E:S ratio showed that laminin was evenly distributed in the epithelia relative to the stroma of benign tissue (Fig. 5D). Interestingly, laminin had a higher E:S ratio in non-STIC tissue versus benign tissue, but no difference was observed in STIC tissue compared to benign (Fig. 5D) or between matched non-STIC and STIC regions (Supplementary Figure 5).

**Figure 5.**
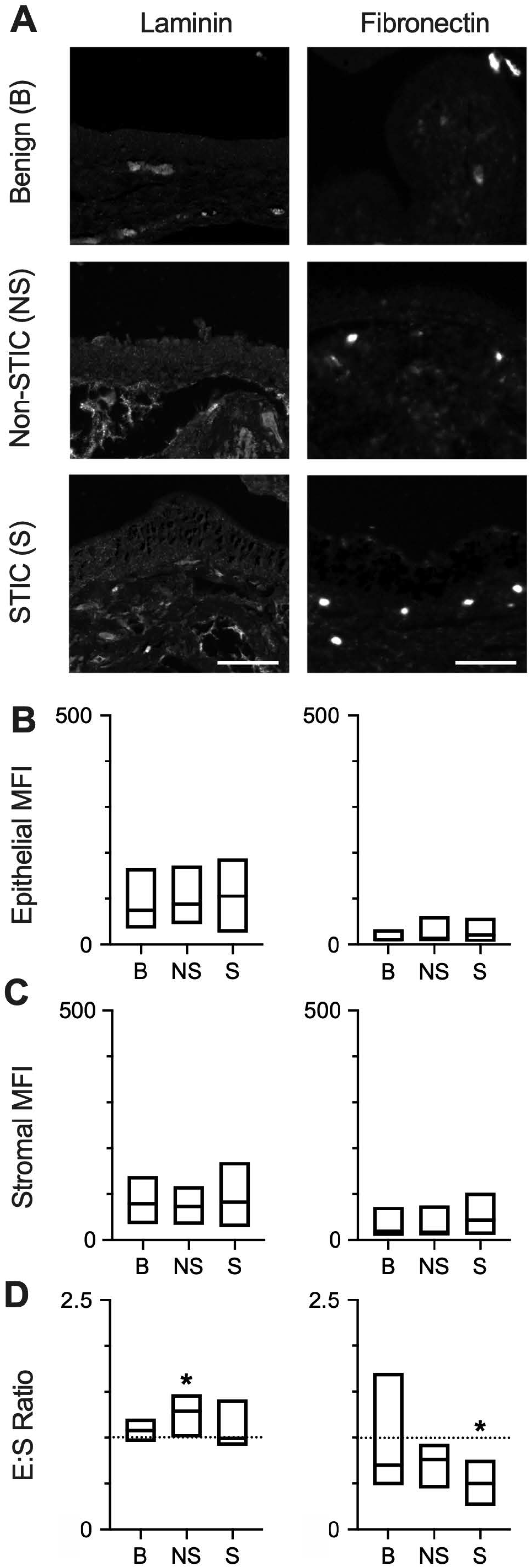
Differential levels of glycoproteins in STICs. (A) Representative multispectral staining of laminins and fibronectin in benign (B), non-STIC (NS), and STIC (S) regions. (B-D) Quantification of (B) epithelial MFI, (C) stromal MFI, and (D) epithelial:stromal (E:S) ratio (of laminins and fibronectin in B, NS, or S regions. Data are presented as box plots indicating median and range, dashed line demonstrates E:S = 1. * indicates p< 0.05 for S compared to B by Kruskal-Wallis test and Dunn’s multiple comparison post-test. For laminins n=10 (B), 8 (NS), 7 (S); for fibronectin n=10 (B), 11 (NS), 8 (S). Scale bar = 200 μm.

The presence of fibronectin was observed in multispectral staining of epithelial and stromal regions of benign fallopian tube/fimbriae, non-STIC regions, and STICs (Fig. 2B). Analysis of epithelial regions showed that levels of fibronectin were unchanged in all comparisons (Fig. 5A-B, Supplementary Figure 5). In the stroma, the levels of fibronectin did not significantly differ when comparing benign tissue to either non-STIC tissue or STIC tissue (Fig. 5C). However, stromal levels of fibronectin were significantly higher in STIC regions compared to non-STIC regions of patient-matched samples (Supplementary Figure 5). Calculation of the E:S ratio demonstrated wide variation in fibronectin distribution across samples, but fibronectin was generally higher in the stroma (median E:S ratio ∼0.6; Fig. 5D). The E:S ratio was significantly lower in STIC tissue compared to benign (Fig. 5D) or compared to non-STIC regions of patient-matched tissue (Supplementary Figure 5). Given the increased levels of stromal fibronectin that were observed in patient-matched samples, it is likely that the relative enrichment of fibronectin in the stromal region during early disease is due to increased stromal fibronectin rather than loss of epithelial fibronectin.

## DISCUSSION

The ECM has been shown to be altered in early tumors, leading to effects on cellular behavior that support cancer progression [36, 37]. Therefore, we sought to characterize the ECM microenvironment of the human fallopian tube and examine how the ECM is altered in early STICs. We first conducted a meta-analysis of several global proteomic datasets that included benign fallopian tubes [19] or the histologically normal fallopian tube tissue from patients with metastatic HGSOC [18, 20, 21]. We then characterized the matrisome of a benign human fallopian tube using the mass spectrometry pipeline we previously developed [22]. A comparison of the overlap between these data sets demonstrated a large overlap for core matrisome proteins. This finding suggests that matrisome annotation of global proteomic data sets such as data generated by the National Cancer Institute Clinical Proteomic Tumor Analysis Consortium (CPTAC, https://proteomics.cancer.gov/programs/cptac) could be useful to identify changes in the matrisome during tumor progression. In addition to the HGSOC tumors analyzed here, CPTAC has characterized the proteomes of over 3,000 samples from 12 tumor types using a standardized method.

Importantly, the meta-analysis revealed a large number of matrisome-associated proteins that were not identified in our insoluble-matrix enriched sample. This finding is perhaps not surprising as some of these proteins may be at much higher abundance intra-cellularly than in the matrix. It will be interesting to further fractionate peptide samples, for example using basic reversed phase liquid chromatography [22, 25], to identify proteins present in lower abundance in the insoluble ECM scaffold, such as growth factors or cytokines, to determine which of these factors could regulate the microenvironment. While a recent study of the ampullar region of the fallopian tube demonstrated minimal changes between the luteal and follicular phase for six ECM analytes [16], there remains the potential for patient-to-patient variability with respect to age and hormonal status (i.e., pre- vs. post-menopausal, hormonal interventions, and menstrual cycle stage). To clarify the impact of these variables on the matrisome, a larger bank of benign fallopian tube samples could be collected and analyzed.

Comparison of the matrisome from the insoluble matrix-enriched sample to the global proteomic data identified seven additional proteins: collagens (COL2A1, COL4A5, COL16A1), glycoproteins (elastin, α chain of laminin 5), annexin A2 and PAI1. mRNA expression of all but COL2A1 have been reported in benign human fallopian tubes (www.proteinatlas.org; [38]). Interestingly, several of these proteins have been linked to later stage events in HGSOC metastasis. Annexin A2 is found both intracellularly and in the stroma of metastatic HGSOC at higher levels than normal ovarian tissue, with potential roles in metastatic adhesion and invasion into the peritoneum [39]. PAI-1 levels have been shown to correlate with peritoneal metastasis and a worse prognosis in HGSOC [40]. LAMA5 has been shown to be expressed in endometrioid ovarian cancer [41] and ovarian clear cell carcinoma [42], but to our knowledge has not be described in HGSOC. Decreased COL2A1 expression [43] and elevated COL16A1 expression [44, 45] have been correlated to poor prognosis in HGSOC. Finally, one study demonstrated a decrease in tropoelastin expression in ovarian tumors relative to normal ovarian tissue [46]. We would emphasize that many of these studies utilized ovarian tissue as a comparison; therefore, much remains to be learned about the role of these proteins in the normal fallopian tube and in the early development of HGSOC tumors.

We utilized the overlap between global proteomics and our matrisome-specific analysis to identify targets to characterize by multispectral imaging in benign tissue and STICs. Our results demonstrated that there were no significant differences in epithelial or stromal ECM levels between benign tissue and non-STIC regions from patients with STICs. However, it is unclear if this conclusion would hold as the number and size of STICs increased or after the onset of metastasis and the accumulation of ascites. Indeed, prior work from our lab has demonstrated that soluble factors in the ascites are capable of increasing ECM deposition in tissue that is distal from the tumor mass [47]. Therefore, further characterization of the STIC matrisome should rely on tissues acquired in the absence of tumors. Our results also demonstrated substantial variation in the levels of several ECM components, particularly in STICs. We therefore relied on the ability to use patient-matched STIC and non-STIC regions to control for variables such as age and hormonal status. These comparisons identified three variations: decreased versican in the STIC epithelium, increased fibronectin in the STIC stroma, and a trend towards increased ECM in the stroma relative to the epithelium in STICs for type III collagen, type IV collagen, perlecan, versican, and fibronectin.

Increased stromal matrix is a common feature of many tumors [36], and is commonly associated with the production of increased matrix by cancer-associated fibroblasts (CAFs, [48]). To our knowledge, the CAF phenotype has not been observed in early STICs; additionally, the stroma that was characterized in our studies was immediately proximal to the epithelial layer and may result from changes in ECM expression and remodeling within the fallopian tube epithelium. The observation of increased fibronectin in the stroma of these precursor lesions is consistent with findings from matrisome characterization of metastatic HGSOC, where fibronectin showed the greatest fold change when comparing omental tumors and omentum from Stage I/II disease that has not spread throughout the peritoneum [49]. A potential role for fibronectin has been reported in the transition from STIC to ovarian tumor [50] and mouse oviductal epithelial cells have been shown to adhere to fibronectin [51].

Intriguingly, our results demonstrated a decrease in versican in the fimbriae epithelium while there were minimal changes in stromal levels. In contrast, prior work has demonstrated an increase in versican in many tumor types (including HGSOC [11]), with elevated levels a prognostic indicator in some solid tumors [52]. In general, reduction of versican has been shown to decrease cell proliferation and increase apoptosis [53, 54]; responses that would be associated with a pro-tumorigenic role for versican. While further work will be needed to elucidate the potential effects of decreased versican on STIC formation, the literature does suggest two possibilities. First, versican has been described as ‘anti-adhesive’ in that it can interfere with binding to fibronectin [55-57]. Therefore, the combination of increased stromal fibronectin with decreased epithelial versican may alter cellular organization and set the stage for epithelial cells to develop the multi-layered structure characteristic of a STIC. Second, versican can impact expression of both E-cadherin and N-cadherin [58, 59] which could impact cellular organization of the STIC.

In conclusion, out study resulted in the first matrisome-specific characterization of the human fallopian tube. Using this information and an imaging analysis method appropriate for small, clinical samples, we identified changes in the epithelial:stromal abundance of several ECM proteins, increased stromal fibronectin, and decreased epithelial versican in STICs. These results suggest that similar approaches will be useful to identify changes in the ECM of other early tumors that have limited sample material.

## DECLARATIONS

### Availability of data and materials

The mass spectrometry dataset generated during this study has been deposited to the ProteomeXchange Consortium via the PRIDE partner repository (PXD023707). The additional data generated during this study are available from the corresponding author on reasonable request.

### Competing interests

The authors declare they have no competing interests.

### Funding Statement

Funding was provided by a pilot award from NIH P30CA014520-46 (KSM) and NIH R01CA232517 (PKK, KSM, AN). Proteomics services were provided by the UIC Research Resources Center Mass spectrometry Core which was established in part by a grant from The Searle Funds at the Chicago Community Trust to the Chicago Biomedical Consortium. Bioinformatic analyses were performed by the UIC Research Informatics Core, supported in part by the National Center for Advancing Translational Sciences (NCATS, Grant UL1TR002003). INT is the recipient of a UIC Honors College Research Grant and an award from the LAS Undergraduate Research Initiative (LASURI) at UIC. The funders had no role in study design, decision to publish, or preparation of the manuscript.

### Author contributions

CR, AN, KSM, and PKK designed the study, PW and SM identified clinically appropriate samples and helped with data interpretation, CG, IT, and AN conducted the analysis and experiments for Fig 1 and prepared Fig. 1 and associated supplements, CR conducted the experiments for Fig 2-5, MRV and AK did the quantification in Fig. 3-5 and associated supplements, MRV prepared Fig. 2-5 and associated supplements, CR, CG, MRV, AN, KSM, and PKK wrote the manuscript. All authors have read and approved the final manuscript.

### Ethics approval and consent to participate

This study was reviewed by the University of Wisconsin IRB and declared to be not humans subject research as all materials were deidentified.

### Consent for publication

Not applicable.

## Acknowledgements

We thank the University of Wisconsin Carbone Cancer Center Experimental Pathology Laboratory and the University of Wisconsin Optical Imaging Core supported by NIH5P30CA014520. We also would like to thank Dr. Hui Chen and Dr. Sunil Hwang from the Mass Spectrometry Core facility at the University of Illinois at Chicago and Dr. George Chlipala from the Research Informatics Core facility at the University of Illinois at Chicago for their technical assistance. We thank the members of the Kreeger, Masters, and Naba laboratories for helpful discussions with this manuscript.

## LIST OF ABBREVIATIONS

ACN: acetonitrile
ECM: extracellular matrix
E:S: ratio of epithelial to stromal MFI
FA: formic acid
FT: fallopian tube
FDR: false discovery rate
GAGs: glycosaminoglycans
HA: hyaluronic acid
HGSOC: high grade serous ovarian cancer
HSPG2: perlecan
MFI: mean fluorescent intensity
ROI: region of interest
SEE-FIM: sectioning and extensively examining the fimbriated end
STIC: serous tubal intraepithelial carcinoma
TBST: tris-buffered saline with Tween
TFA: trifluoroacetic acid
TP53: tumor protein P53
VCAN: versican

## SUPPLEMENTARY TABLES

**Supplementary Table 1:** Meta-analysis of global proteomic data set on protein composition of human fallopian tubes

**Supplementary Table 2:** De novo human fallopian tubes proteomic dataset with ECM enrichment

**Supplementary Table 3:** Human fallopian tube samples used for multispectral immunohistochemistry

**Supplementary Table 4:** List of antibodies used in multispectral IHC

---

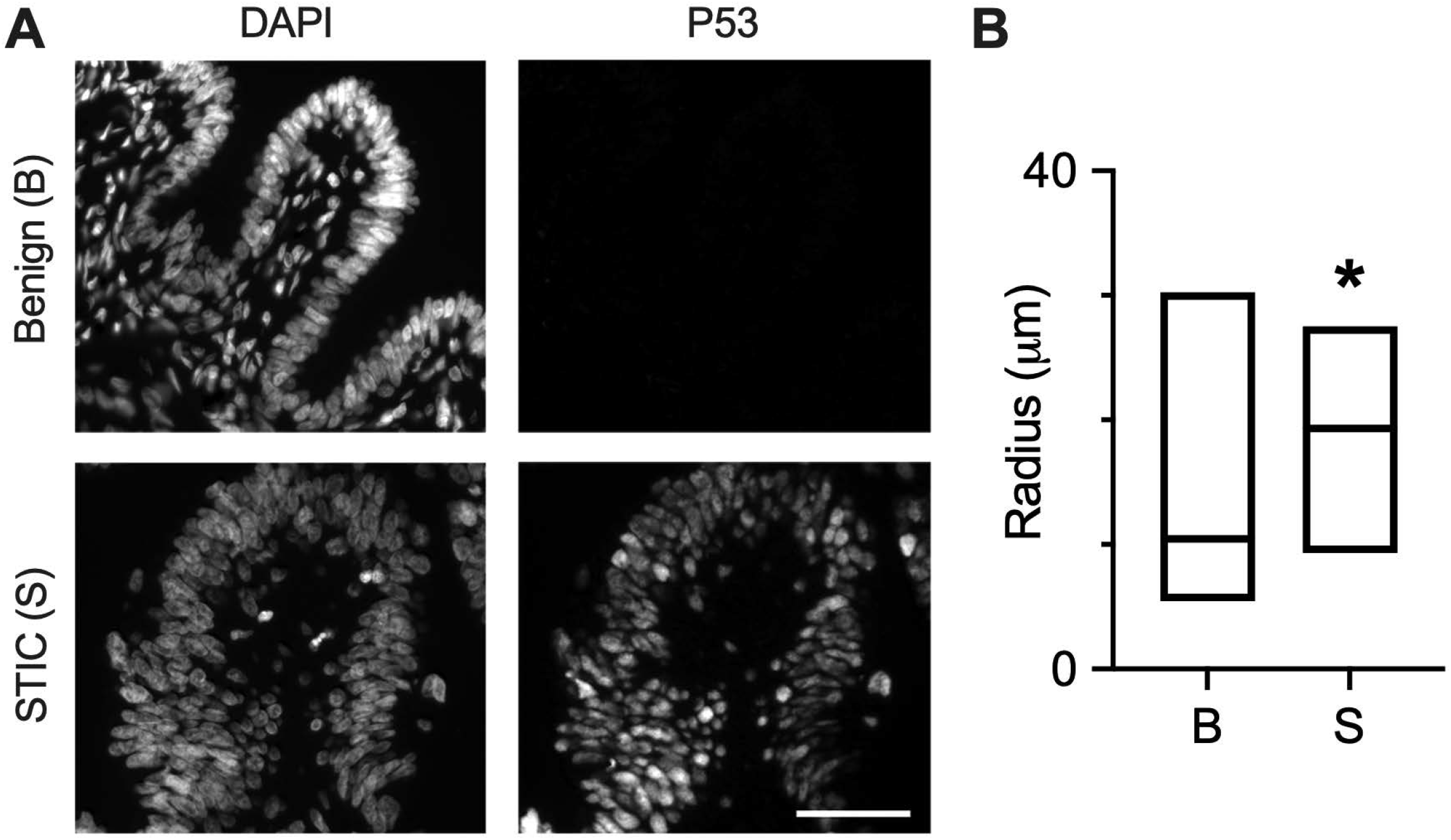

## Notes

### Competing Interest Statement

The authors have declared no competing interest.

