## Supplementary Figures for "Multi-modal profiling of the extracellular matrix of human fallopian tubes and serous tubal Intraepithelial carcinomas"

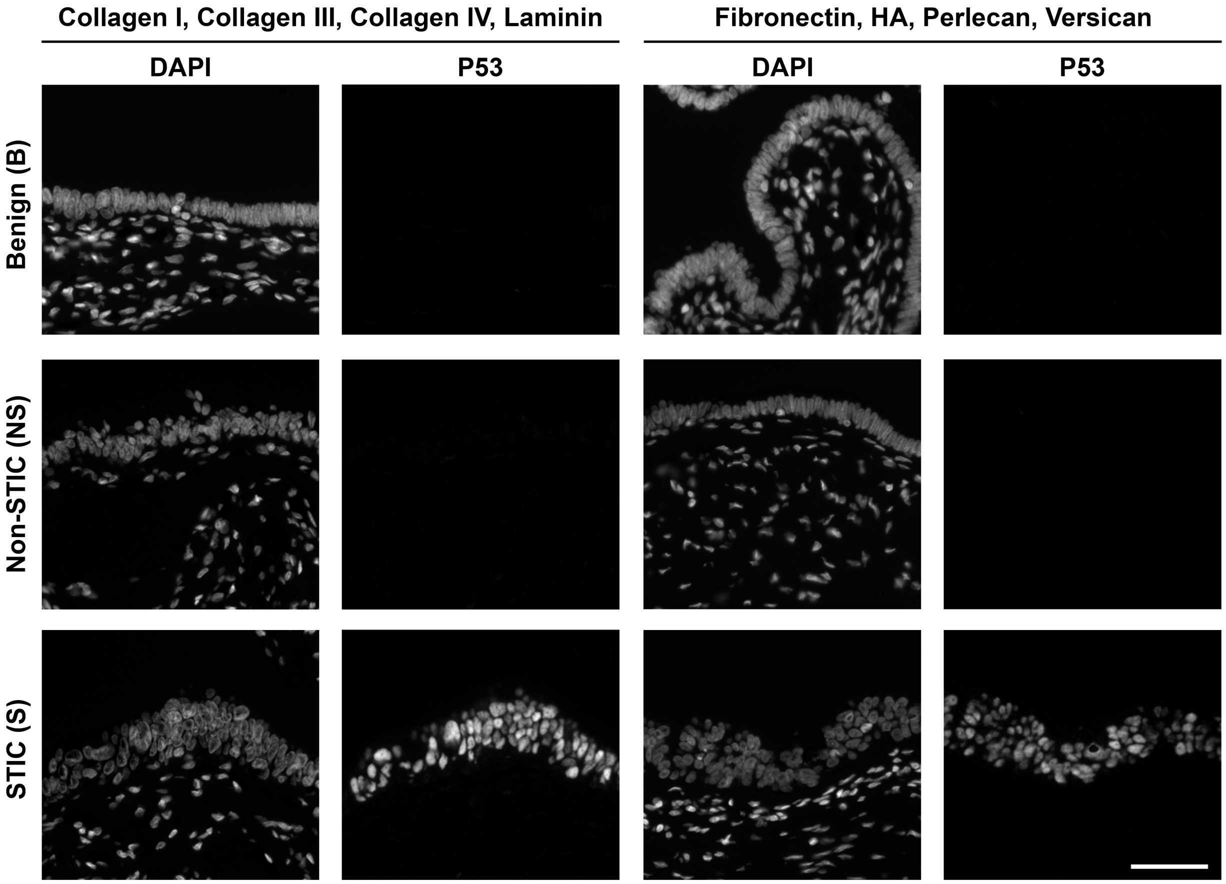


***Supplemental Figure 1.*** *Nuclear and P53 staining of fallopian tube/fimbriae samples.* Nuclear signal (DAPI) and P53 immunostaining of benign (B), non-STIC (NS) and STIC (S) regions; these images correspond to the representative ECM immunostaining images shown in **Fig. 3-5**. Scale bar = 200 $\mu$m.

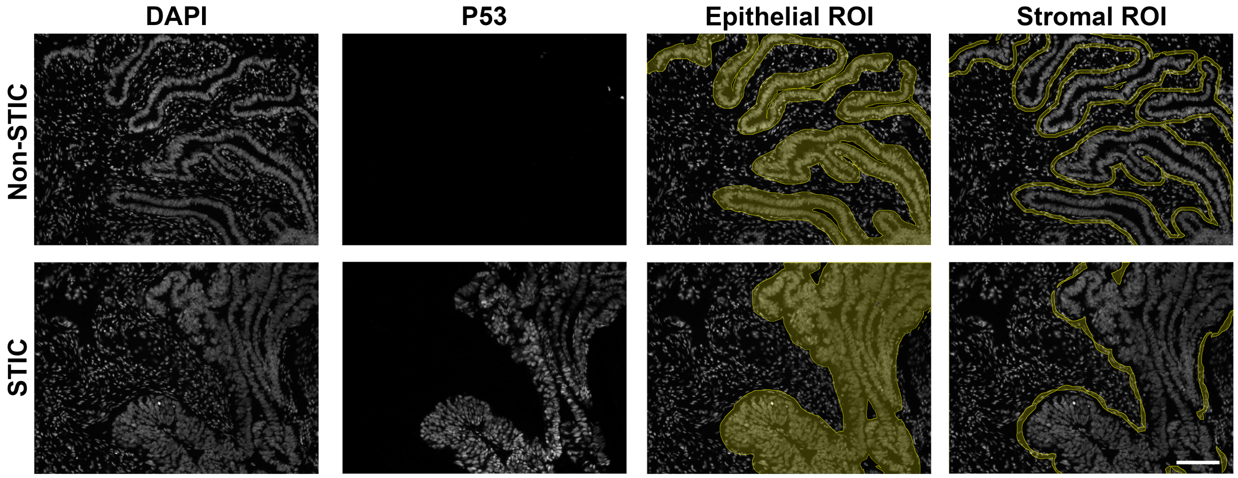


***Supplemental Figure 2.*** *ECM quantification from multispectral imaging.* Nuclear (DAPI) and P53 staining of Non-STIC and STIC regions are shown to the left. These images were used to define the epithelial and stromal regions in FIJI (shown on right). Briefly, the epithelial of non-STIC regions was manually outlined using the DAPI channel image. Separately, the nuclei in the complete image were selected using the threshold tool in the DAPI channel. A new selection featuring just the nuclei within the epithelial region was created from the epithelial selection and nuclei selection using the 'AND' function in the ROI manager. This region was expanded by 25 $\mu$m to define the epithelial ROI. The stromal region of the non-STIC was defined using the 5 $\mu$m region beyond the epithelial ROI. STIC regions were defined similarly using the P53 channel expanded by 30 $\mu$m to define the epithelial ROI and an additional 5 $\mu$m to define the stromal ROI.

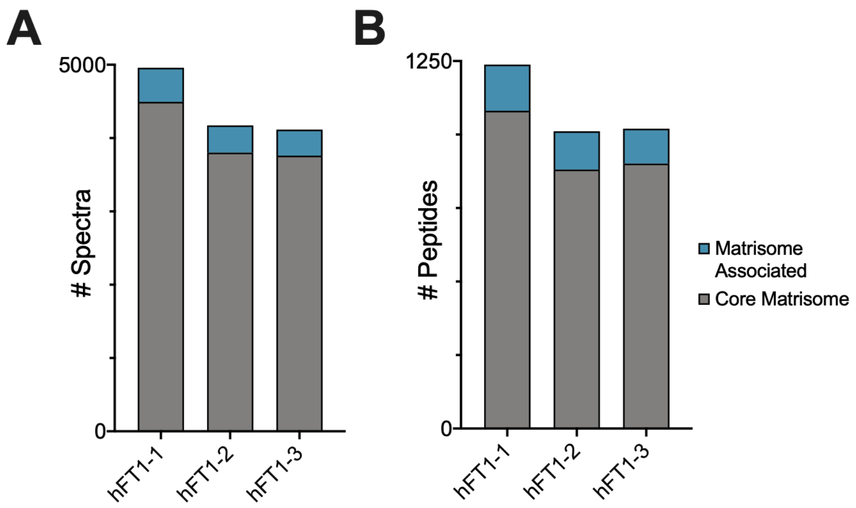


***Supplemental Figure 3.*** *The analytical pipeline demonstrated high reproducibility. The three technical replicates (hFT1-1, hFT1-2, hFT1-3) led to the detection and identification of similar number of spectra (left panel) and peptides (right panel) (see* ***Supplemental Tables 2B and 2C****)*

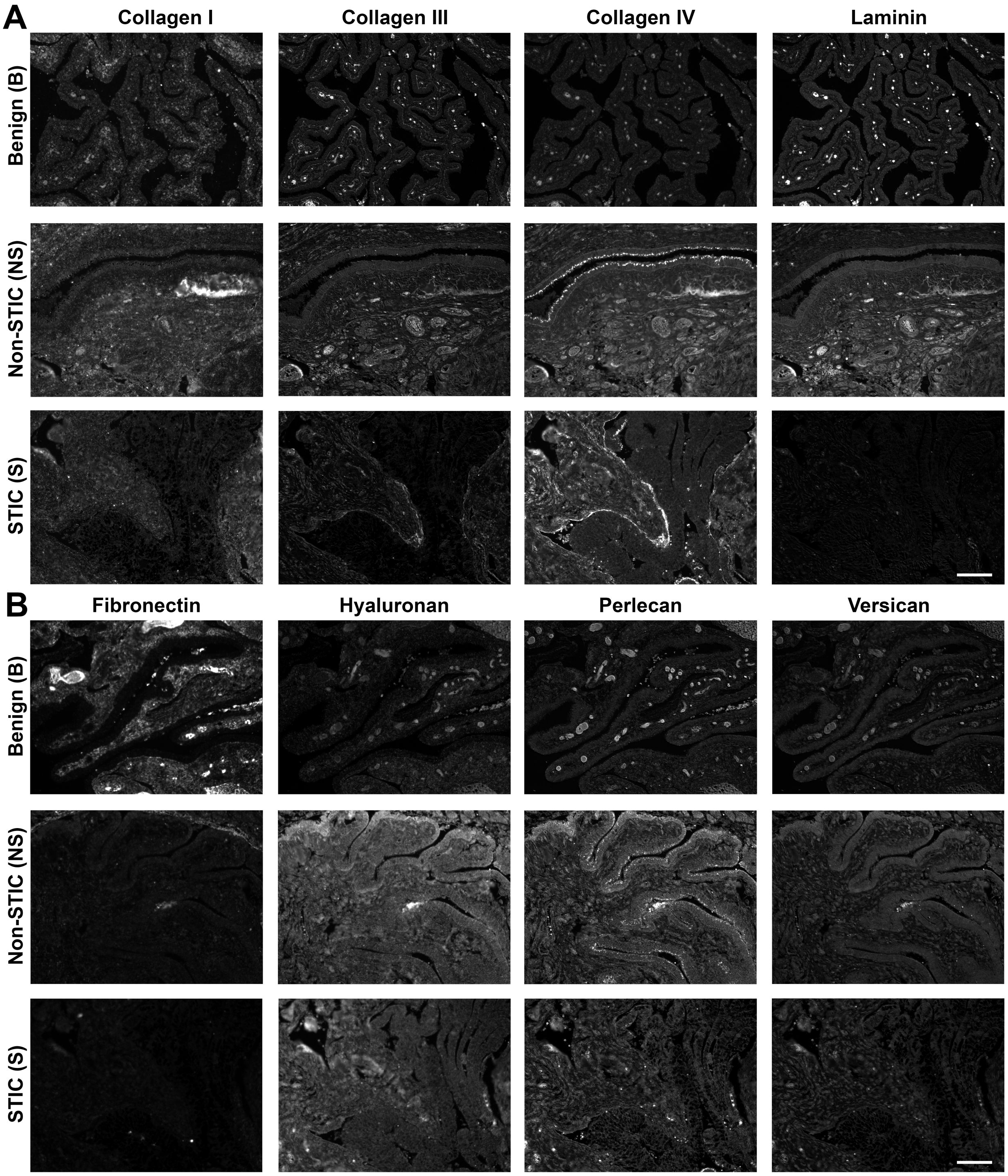


***Supplemental Figure 4.*** *Immunofluorescent images of ECM components in fallopian tube/fimbriae.* Collagen I, collagen III, collagen IV, pan-laminin, fibronectin, hyaluronan, perlecan, and versican immunofluorescent images of benign (B), non-STIC (NS) and STIC (S) regions; these images correspond to individual channels of the representative ECM immunofluorescent images shown in **Fig. 2**. Scale bar = 200 μm.

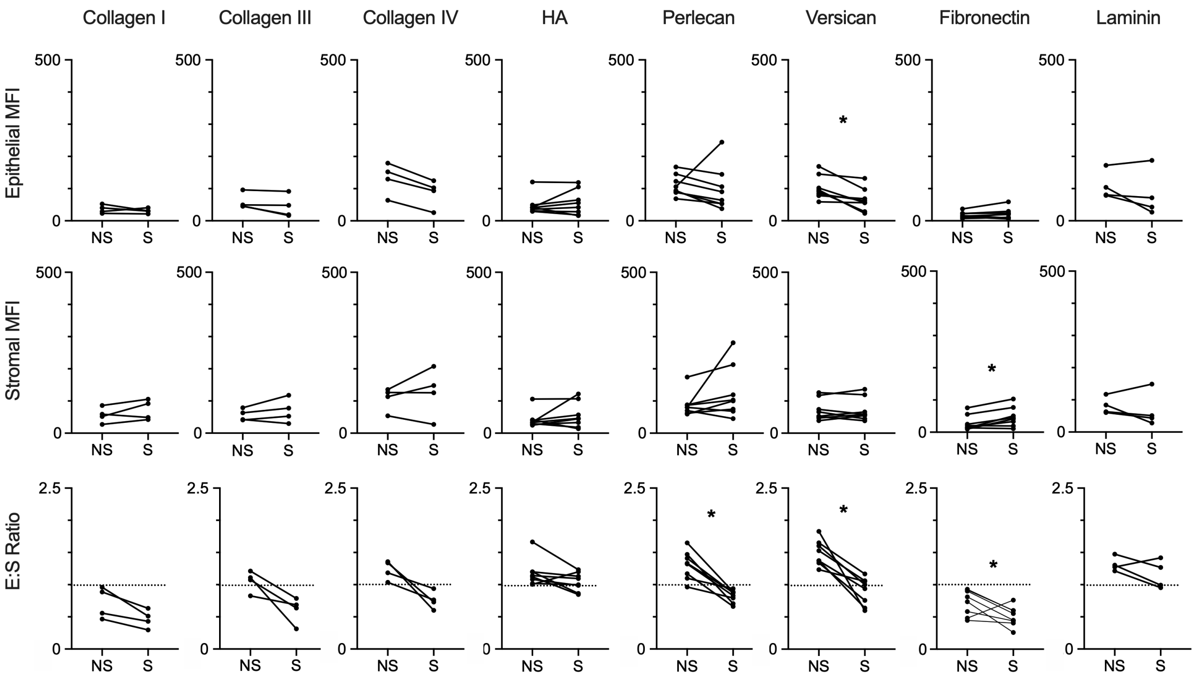


***Supplemental Figure 5.*** *Comparisons of non-STIC (NS) versus STIC (S) regions in patient-matched samples.* Epithelial MFI, Stromal MFI, and E:S ratio of patient-matched regions (indicated by connecting lines). Dashed line demonstrates E:S = 1, * indicates $p<0.05$ by Wilcoxon matched-pair signed rank test. For fibronectin, hyaluronan, perlecan, and versican, n=8 patients. For collagen I, collagen III, collagen IV, and laminin, n=4 patients
